# PyRodentTracks: flexible computer vision and RFID based system for multiple rodent tracking and behavioral assessment

**DOI:** 10.1101/2022.01.23.477395

**Authors:** Tony Fong, Braeden Jury, Hao Hu, Timothy H. Murphy

## Abstract

PyRodentTracks (PRT) is a scalable and customizable computer vision and RFID-based system for multiple rodent tracking and behavior assessment that can be set up within minutes in any user-defined arena at minimal cost. PRT is composed of the online Raspberry Pi-based video and RFID acquisition and the subsequent offline analysis tools. The system is capable of tracking up to 6 mice in experiments ranging from minutes to days. PRT maintained a minimum of 88% detections tracked with an overall accuracy >85% when compared to manual validation of videos containing 1-4 mice in a modified home-cage. As expected, chronic recording in home-cage revealed diurnal activity patterns. Moreover, it was observed that novel non-cagemate mice pairs exhibit more similarity in travel trajectory patterns over a 10-minute period in the openfield than cagemates. Therefore, shared features within travel trajectories between animals may be a measure of sociability that has not been previously reported. Moreover, PRT can interface with open-source packages such as Deeplabcut and Traja for pose estimation and travel trajectory analysis, respectively. In combination with Traja, PRT resolved motor deficits exhibited in stroke animals. Overall, we present an affordable, open-sourced, and customizable/scalable rodent-specific behavior recording and analysis system.

**Statement of Significance:** An affordable, customizable, and easy-to-use open-source rodent tracking system is described. To tackle the increasingly complex questions in neuroscience, researchers need a flexible system to track rodents of different coat colors in various complex experimental paradigms. The majority of current tools, commercial or otherwise, can only be fully automated to track multiple animals of the same type in a single defined environment and are not easily setup within custom arenas or cages. Moreover, many tools are not only expensive but are also difficult to set up and use, often requiring users to have extensive hardware and software knowledge. In contrast, PRT is easy to install and can be adapted to track rodents of any coat color in any user-defined environment with few restrictions. We believe that PRT will be an invaluable tool for researchers that are quantifying behavior in identified animals.

## 1. Introduction

Paradigms have been developed for identifying abnormal behavioral phenotypes in animal models of neuropsychiatric disorders and drug screening in pharmacological studies (Sourioux et al., 2018). Traditionally, these approaches rely on manual phenotyping which is time, labor, and skill intensive. At the same time, results are not only prone to investigator bias and handling effects on animals (Ohayon et al., 2013), but also random errors dependent on the evaluator. As expected, results from traditional paradigms are usually high in inter-experiment variability and often difficult to reproduce (Kafkafi et al., 2018). Moreover, these paradigms are usually session-based and only present a single snapshot of possible existing animal behavior phenotypes (Bains et al., 2018). There is an increasing need for behavior assays to be fully automated and capable of long-term assessments to capture a more complete sample of behaviors.

Here, this report presents PyRodent Tracks: an affordable, open-sourced, easy-to-set- up, and customizable/scalable behavior recording software and hardware. The system is capable of recording and tracking multiple mice of varied coat colors for extended periods of time in any user-defined environment. Video and RFID recordings were done utilizing a Raspberry Pi (RPi) 3B+/4 micro-computer. In the offline pipeline, mice are first detected and tracked using the You Only Look Once version 4 (Yolov4) algorithm coupled with a modified version of the Simple Online and Realtime Tracking (SORT) for mice tracking, respectively (Bewley et al., 2016; Bochkovskiy et al., 2020).

Subsequent posture detections and travel trajectory analysis can also be done by interfacing with multi-animal DeepLabCut (Lauer et al., 2021) and Traja (Shenk et al., 2020), respectively. PRT home-cage recording used in our home-cage study contains 6 RFID readers and only costs approximately 520 USD per mouse cage. The RFID reader numbers and locations can be adjusted to other home-cage variations or recording environments to to fit a specific investigator needs. To demonstrate flexibility and scalability, tracking of six mice in open-field (781 USD) and three white coat color coated mice in a three chambers arena (428 USD) against a low contrast background were also performed.

## 2. Materials and Methods

### PRT Recording Setup

All components are connected to and controlled by custom software running on a RPi 3B+/4 micro-computer running the Raspbian Buster (https://www.raspberrypi.org) as seen in Fig 1A. During each recording, frames are written to an H264 video file while the timestamp of each frame is collected in a separate csv file. RFID reader output is also recorded throughout the session. Hardware installation of the recording system follows the rule of plug-and-play with minimal hardware modification needed and few restrictions on the rodent arena employed (Supplemental Video 1 and 2). Depending on the resolution, a maximum frame rate of 90 fps at 640 x 480 can be achieved using the RPi V1 camera. For further information on the possible frame rate and resolution combination please refer to the official picamera documentation (https://picamera.readthedocs.io/en/release-1.12/fov.html).

**Figure 1.**
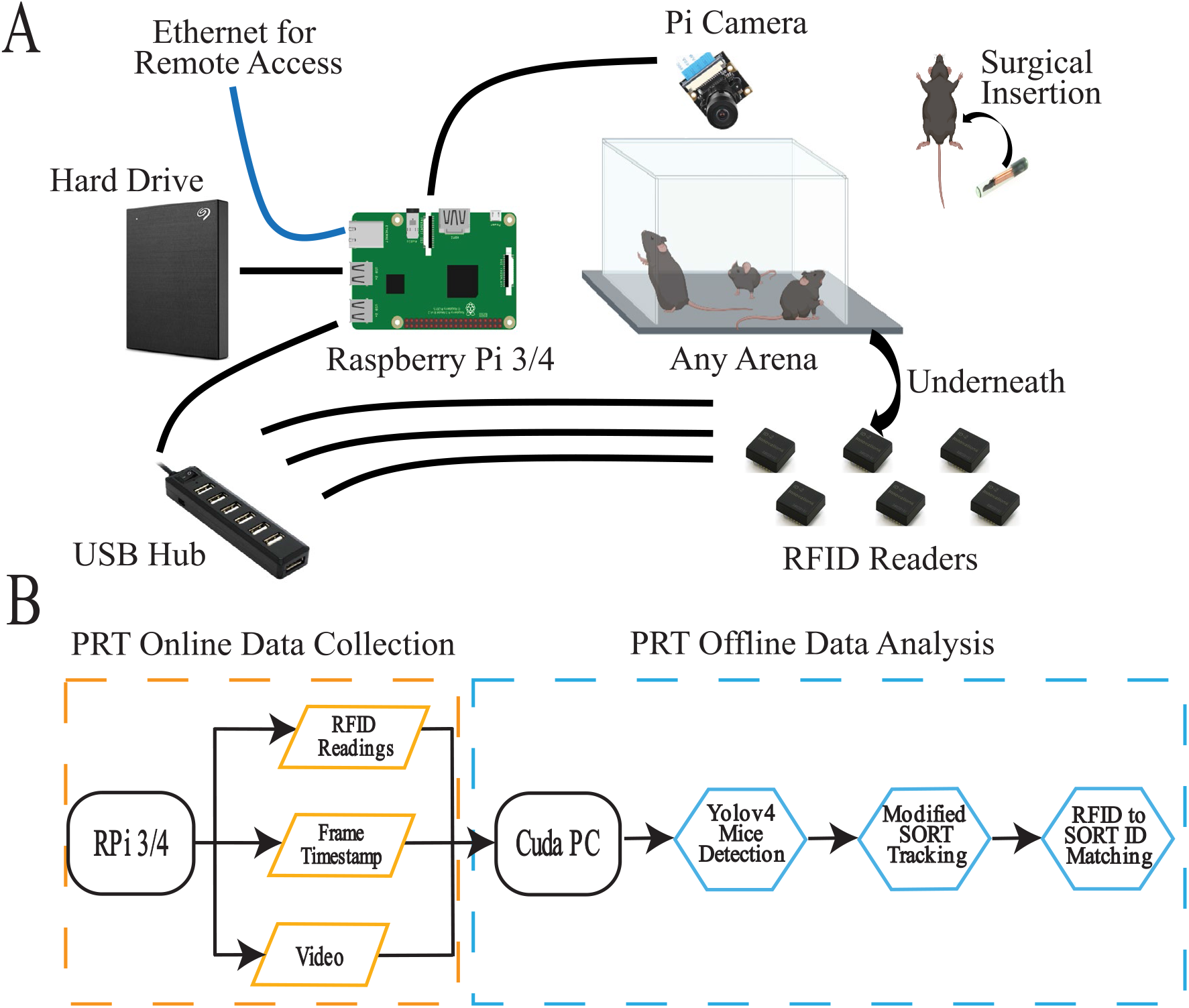
PRT data collection and offline data analysis. **A)** The essential hardware and general recording setup of PRT. The recording can be done in any arena. Main components include a RPi 3B+/4 connected to an overview Pi Camera of the arena, a powered USB hub to relay the RFID readers to the Pi, a hard drive for storage, and RFID readers underneath the arena. Mice being recorded have been surgically implanted with an RFID tag. When connected to ethernet, the system can be remotely accessed and controlled. **B)** PRT RPi online data collection and offline analysis pipeline. Data is collected on a RPi 3/4 which records to a video, while simultaneously recording the timestamp of each frame and RFID readings from RFID readers. Data collected can be analyzed offline using a Colab notebook or a cuda-capable PC.

In brief, a pi-camera is situated above the rodent arena with an unobstructed view (any RPi compatible camera can be used). RFID readers are connected to RPi through a self-powered USB hub and situated underneath the arena housing the RFID tag implanted rodents. The number of RFID tag readers used can be adjusted to arenas of any dimension as long as they are spaced 12 cm apart to minimize RFID interference. Currently, up to 9 readers have been tested. Connection to ethernet would allow remote access and control of the RPi. All essential parts are listed in Table 1. We also recommend installing a supplemental RPi cooling solution.

**Table 1.** Essential components for building the PRT online recording system.

| Component | Amount Required | Supplier | Part Number | Notes |
| --- | --- | --- | --- | --- |
| Raspberry Pi 3B/4 | 1 | Newark | RASPBERRYPI3-MODB-1GB/RASPBERRYPI4-MODB-4GB | Either model could be used |
| RFID Reader ID20-LA | Depends on user | Sparkfun | SEN-11828 | As many as the user wants as long as there are spaced a minimum of 12 cm apart |
| RFID Reader Breakout | Depends on user | Sparkfun | SEN-13030 | Matching to the number of RFID Reader ID20-LA |
| USB Mini B Cable | Depends on user | AmazonBasics | N/A | Or equivalent. |
| RFID Glass Capsules | 1 | Sparkfun | SEN-09416 | One needed per animal to be tested. |
| 16GB Micro-SD Card | 1 | Scan | 44082 | Or equivalent. |
| Pi Camera(F/G) | 1 | Waveshare | 10299/10344 | Or any other Pi compatible camera |
| Portable Hard Drive | 1 | Seagate | STKC4000400 | Or equivalent. |
| RPi Cooler | 1 | Smraza | SW49 | Optional but recommended. Other equivalent options could be used |
| Thermal Paste | 1 | Noctua | NT-H1 | Optional but recommended. Other equivalent options could be used |
| PLA Filament, 1.75mm Black | Depends on user | MakerBot | MP05775 | Optional. To print custom 3D parts for home-cage design or RFID reader holder. Other equivalent options could be used |

### PRT Offline Analysis Pipeline

The PRT offline analysis pipeline runs on a CUDA-installed computer in an Anaconda environment (https://www.anaconda.com). The pipeline mainly contains three components as seen in Fig 1B and Fig 2A: 1) Yolov4 for mouse detections (Bochkovskiy et al., 2020), 2) modified SORT for mice tracking(Bewley et al., 2016), 3) in house generated RFID to SORT ID matching for identity confirmation. Mice were first detected by a TensorFlow 2 framework of Yolov4 recognized by the original creators (Hùng, 2021). All mouse detections, as bounding boxes, from Yolov4 were then assigned an initial underlying ID by a modified SORT algorithm which uses the Kalman filter to predict and track each mouse. As seen in Fig 2B, the two main features of the modified SORT algorithm include Kalman filter predictions and SORT-ID reassociation to enhance detection and tracking performance. As the maximum number of mice is known, the Kalman filter can therefore predict mouse position when Yolov4 fails, such as in cases of visual occlusions or detections being in close proximity. Similarly, new false positive SORT-IDs can be associated with a previous ID that disappeared. At times, new SORT-IDs will be generated for the same mouse that is already tracked, discarding a previous tracked ID. The generation of these new false SORT-IDs is due to the irregular travel patterns taken by mice which remains difficult to completely be predicted by SORT’s underlying Kalman filter. With the modifications combined, the modified SORT algorithm provides highly accurate tracking of individual animals during clustering scenarios (Supplemental Video 3).

**Figure 2.**
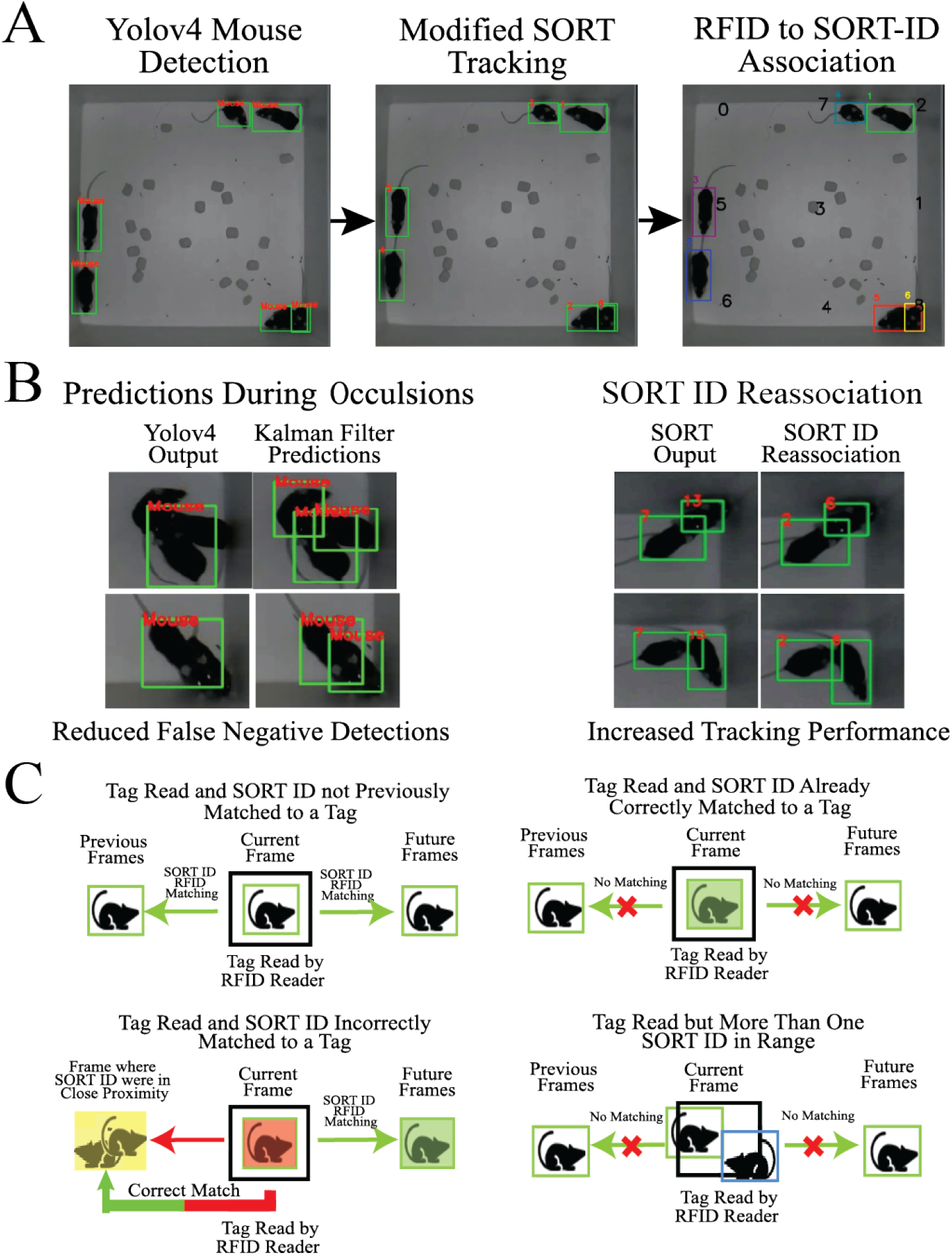
Overview of the PRT offline analysis pipeline. **A)** Mice are first detected by Yolov4 and tracked using a modified SORT tracker. Then the SORT IDs generated from the SORT tracker are temporally matched to RFID tags read by the RFID reader. **B)** Main features of the modified SORT tracker. In cases of Yolov4 detection failures such as visual occlusions or close proximity of detections, the Kalman filter will still output predictions of mice. The second feature is the re-association of false-positive SORT-ID to an old SORT-ID that disappeared to increase tracking performance. **C)** The possible scenarios and SORT-ID to RFID matching outcomes when an RFID tag is read by an RFID tag reader.

In the last stage of PRT offline analysis, the SORT-IDs generated are matched with RFID readings. The general overview of the matching process is illustrated in Fig 2C. RFID reader locations are user defined regions of interest on the video image captured. When a tag is read by an RFID reader, there are four possible scenarios which then, in turn, can lead to three possible outcomes for the matching process: 1)SORT-ID to RFID matching in all previous and future frames, 2) No Matching, and 3) Matching of future frames and correction of previous frames to a point of occlusion. Moreover, both centroid distance and intersection over union (IOU) were used to determine if a SORT-ID detection is in the range of an RFID reader. In the first scenario, the SORT ID has not previously been matched to a tag and therefore, it will be matched with the tag in all previous and future frames. In the second scenario, the SORT-ID has already been correctly matched to the tag being read, so no matching would occur. In the third scenario, there is more than one SORT-ID (i.e. more than one mouse) in the range of the RFID reader. To ensure clean RFID readings and proper matching, again no matching would occur. In the final scenario, the SORT-ID is incorrectly matched to a tag. Therefore, all SORT-ID to RFID matches in future frames and previous frames up to a point where it was in close proximity with another SORT-ID, a situation where identity swaps are likely to occur due the use of IOU to track mice, are corrected.

### Yolov4 Training and Weight Generation

Weights for Yolov4 detection were trained in its original Darknet(Bochkovskiy et al., 2020) implementation through transfer learning using pre-trained weights as the performance has not been fully reproduced on the TensorFlow 2 framework (Hùng, 2021). Trained weights were then converted to Tensorflow weights for the detection to run natively in Python. For the home-cage experiments, weights were trained using over 2000 random, distinctive images from multiple experiment sessions observing 3-4 mice. Similarly, weights for the open-field and three-chamber experiments were trained on 300 random, distinctive images. A minimum mean average precision (mAP) value >99.5% was achieved on the test dataset for all weights (Supplement Fig 1).

### Home-Cage Recording Setup AND PRT Pipeline Line Adjustments

To perform chronic recordings in a home-cage, a shoebox-sized mouse home cage (19 x 29 x 12.7 cm)was modified to hold a water bottle holster, a housing area, and its connecting tunnel as seen in Fig 3A. Moreover, a single fisheye lens camera and IR lights were fitted to the top of the cage. A sixth reader is attached to the tunnel leading to the mouse housing area. A custom-cut acrylic sheet is also placed between the Pi camera with its related wirings and the cage main body.

**Figure 3.**
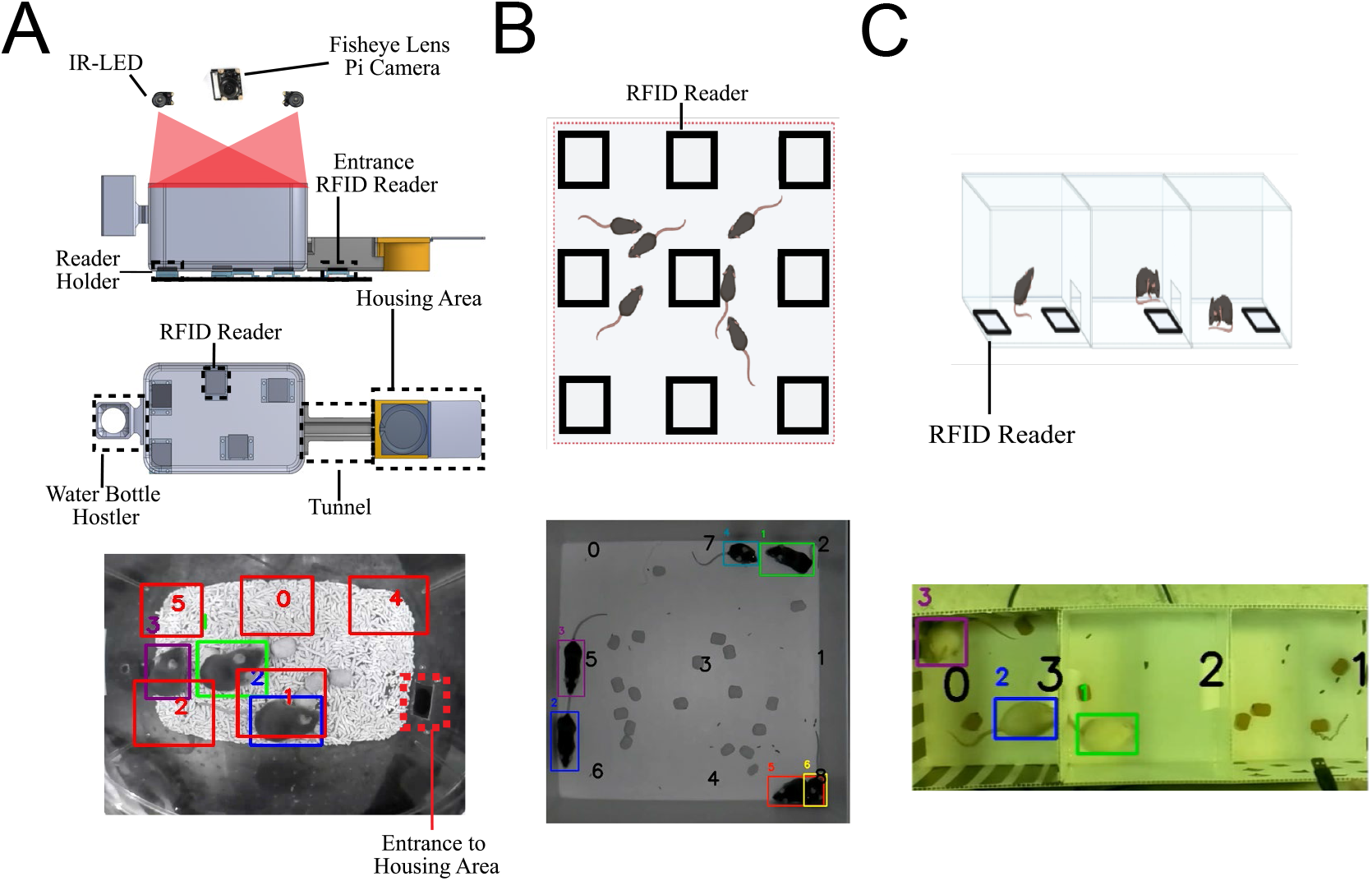
Setup of the Custom home-cage. **A)** The custom home-cage for chronic behavior recording. A fish-eye lens Pi camera and two IR lights are situated at the top of the cage separated by an acrylic sheet. The cage sits on top of the RFID readers which are held by holders. A tunnel is connected to the main cage from one end to the housing area, while the other end holds a water bottle hostler. All home-cage videos were recorded at a resolution of 512 x 400 that was still sufficient to resolve the animals. RFID reader locations are denoted by numbers 0-5 with red rectangles. **B)** PRT recording under IR light and analysis in an open- field arena with 6 black coat-colored coated mice. Video was recorded at 960×960 at 40 fps. **C)** PRT recording under natural light in a three-chamber sociability arena with 3 white coat color coated mice. Video was recorded at 640×480 at 60 fps. RFID readers are denoted by black open box with numbers in the center. Each color and corresponding ID represent RFID tracking of an individual mouse.

For performance validation, fifteen 10-minute videos were recorded containing 1-4 mice. For chronic recordings, mice were left in the home-cage for three days. Food pellets were distributed on the floor of the main cage. Recordings were temporarily interrupted for 15 minutes every 2 days for cage cleaning.

Additional modifications to the offline tracking pipeline were made to accommodate home-cage recordings : 1)all matching processes from cage RFID readers stop at a certain distance to the housing entrance area as seen in Fig 3C 2) SORT-ID to RFID matching correction, as mentioned above. All corrections were traced back into the closest frame of SORT-ID proximity or at a certain distance to the entrance area. The final modification was the use of an entrance RFID reader for SORT-ID to RFID matching. The reasons for these modifications are that the entrance will provide scenarios where a new mouse may appear, a tracked mouse may disappear, or both may occur.

### Motion Detection

To identify and segregate keyframes of interest, i.e., frames containing mice with active movements as opposed to where mice were idle/resting, a simple motion detector is built into the PRT pipeline. Background subtraction against an accumulated average between current and previous frames was used to detect consecutive frames of motion. Specifically, a Gaussian filter was applied to average pixel intensities across a region of 21 x 21. An absolute value between the current frame and the accumulated weights of previous frames was calculated to yield contours representing regions of motion. Specific settings of the motion detector can also be adjusted within the offline tracking pipeline.

### Open-Field and Three Chamber Recording

Nine RFID readers were placed underneath an open-field (32 x 32 cm) as illustrated in Fig 3B. When recording, the open-field arena was placed in a chamber covered by black- out curtains. Videos were recorded at a resolution of 960 x 960 under IR light using a regular lens Pi camera.

For three chamber testing, four RFID readers were placed underneath an arena (20 x 20 cm for each chamber) as illustrated in Fig 3C. Videos were recorded at a resolution of 640 x 480 under natural background lights using a regular lens Pi camera.

### Animals

Male C57BL/6 mice of varied genotypes were used for the behavior recording in the custom home-cage and the open-field arena. We are not reporting the genotype-specific as the experiments were not powered to make comparisons between different animals. Instead, in this study our goal was to evaluate tracking accuracy using surplus animals and will reserve cross-genotype work for future studies. FVB/N mice, also of varied genotype were used for behavior recording in the three-chamber arena. Mice were housed in a conventional facility in plastic cages similar to the home-cage setup and kept under a normal 12 hr light cycle during and prior to behavior recording. In the behavior recording home-cage, 30 ml of pellet bedding and food pellets were provided in the cage. A pellet-type bedding was used for home-cage recording as less clumping would occur and therefore minimize animal RFID tag distance to readers, however other forms of bedding should work providing bedding is not too deep, deep bedding would increase the distance from tags to readers. All procedures were conducted with approval from the University of British Columbia Animal Care Committee and in accordance with guidelines set forth by the Canadian Council for Animal Care.

### RFID capsule implantation

To enable the identification of group-housed mice, animals were implanted with glass RFID capsules (Sparkfun SEN-09416) prior to recording as described in (Bolaños et al., 2017; Murphy et al., 2020). Briefly, animals were anesthetized with isoflurane and given buprenorphine via subcutaneous injection (0.05 mg/kg) for analgesia. Betadine was applied to disinfect the incision site, and a small incision was made in the lower abdomen. A sterile injector (Fofia ZS006) was then used to insert the RFID capsule subcutaneously below the nape of the neck or at the abdominal (abdominal will yield better performance). The incision was sutured, and the animal was removed from anesthesia, allowed to recover, and then returned to its home-cage. Animals were closely monitored for 3 days to ensure healthy recovery and proper placement of the RFID capsule post surgery. Animals were given a minimum of one week to fully recover following surgery before being used for any experiments.

### Stroke Induction

A photothrombotic occlusion was introduced at a target area between the sensory and motor cortex, stereotactic coordinates (1.5; 0.5) mm from bregma, as previously described (Balbi et al., 2021). Briefly, mice fitted with a chronic transcranial window (Murphy et al., 2020) were injected intraperitoneally (0.1 ml per 10 g body weight) with a photosensitive dye solution –Rose Bengal (RB) (R3877-5G, Sigma-Aldrich USA). Two minutes after the injection, a 40mW diode pump solid-state 532 nm laser attenuated to 11mW through a polarizer was turned on at the target area to induce focal ischemia. 10-minute behavior recordings of individual mice were performed at 1-hour pre-stroke, 1-day post-stroke, and 7- day post-stroke in the open field illustrated in Fig 3B. The center area of the arena is defined as 0.5*width and 0.5*length around the center of the open-field arena.

### Manual Validation of Videos

To evaluate the performance of the PRT tracking system, 10-minute videos were recorded at 15 fps at 512 x 400 from the modified home-cage containing 1-4 animals. Two videos were recorded at 30 fps at 960 x 960 in the open-field arena and one video at 30 fps at 640 x 480 in the three-chamber arena. Videos were then evaluated frame for frame by students for appearances of frames containing false positive detections (FP), false negative detections (FN), incorrect SORT-ID and RFID matches. To objectively compare PRT to existing object trackers, the Multiple Object Tracking Accuracy (MOTA) index proposed by Bernardin et. al. (2006) in a similar fashion to that of LMT was used. Specifically, MOTA was calculated by the following equation:

MOTA = 1 - sum(f) (FN (t) + FP(f) + identity error(f)) / sum(f) (number of mice in the ground truth)

f is defined as a frame. FN is a frame where the total number of mice is not detected but it is there; a FP is a frame where the total number of mice detected but not there. An identity error is a frame where the total number of mice labeled with an incorrect RFID tag; and the number of mice in the ground truth is the number of mice that should be tracked in each frame.

### Social Stimulus Test

A test mouse is left in the open-field arena, as described above, with a stimulus mouse cagemate (mouse from the same cage) or a non-cagemate (mouse from a different cage) for 10 minutes. For all trials, the test mouse was first tested with a cagemate and later with a non- cagemate with a 20-minute gap between trials. As the same mouse within a cage was used as the cagemate mouse, tests with the cagemate first prevented any crossover in odor from a non-cagemate mouse. The cage was cleaned with 70% ethanol between trials.

Track pattern difference score is calculated using the dynamic time warping algorithm published by the DTAI Research Group (Meert et al., 2020). Both trajectories were z- normalized before dynamic time warping alignment to calculate a track difference score expressed in euclidean distance. Spatial Proximity (SP) is calculated by the average of all minimal distances of each point on a trajectory to the other and is also expressed in euclidean distance. Distal trajectory pairs were defined as trajectories segments with an SP> 300, whereas proximal trajectory pairs were defined as trajectories segments with an SP< 300. A full illustration of both calculation processes is shown in Fig 7A. Track difference pattern score was expressed as an average for each trial per test animal.

### Traditional Social Interaction Detection

Social interaction (ITC) was calculated by the sum duration of overlap of enlarged (25% area) bounding boxes of detected mice. An ITC episode was defined as the duration between onset and offset of the overlap between mice bounding boxes. Number of ITC episodes were calculated by the count of ITC episodes and the average duration of ITC episodes were also calculated.

### Statistical Analysis

Data are all presented as mean +/- SEM (standard error of the mean). All associated datasets were tested for normality using the Shapiro-Wilks Normality Test (p>0.05).

Statistical significance was determined using either a multivariate regression analysis (MANOVA) followed by a post-hoc ANOVA or repeated measures analysis of variance (RM-ANOVA) followed by paired Student’s t tests as appropriate using R. The level of significance is denoted on the figures as follows: \**p* < 0.05, \*\**p* < 0.01 and \*\*\**p* < 0.001.

## 3. Results

### PRT Detection and Tracking Performance in Home-Cage

Within the methods section, we describe the setup and implementation of PRT for both standard “shoebox-sized” mouse cages and more complex arenas (Fig 3). To evaluate PRT’s performance and reliability 1-4 mice were placed in the home-cage for approximately 10 min recordings at 15 frames per second (Fig 4). Results were compared to manual human frame-by-frame validation. Fig 4A shows the detection reliability of the pipeline. Both the FP and FN detections remain extremely low in all 14 videos analyzed, never exceeding 3% of the total number of mice in manually labeled ground truth. In regards to tracking, performance was mainly evaluated by coverage, MOTA value, and identity error rate as seen in Fig 4B. Coverage is represented as the percentage of mice detected that are matched with an RFID tag and remained above 90% in the majority of videos analyzed. Specifically, the average coverage for one to four animals was 100.0, 97.9, 96.2, and 91.8 percent, respectively. Among the detections matched with an RFID tag, the identity error rate, i.e. the percent of detections with incorrectly matched RFID tags, was on average 0, 3.78, 1.27, and 3.65 percent of all matched detections in videos of one to four animals. Each 10-minute video would contain on average 6 episodes where detections were matched with an incorrect RFID tag (∼45 frames in length). MOTA index is currently one of the main metrics for determining the effectiveness of a multiple object tracker (Bernardin et al.). In the current study, the average MOTA value for videos of one, two, three, and four animals were calculated to be 0.998, 0.945, 0.973, and 0.949, respectively. As a point of reference, the MOTA values for LMT (Chaumont et al., 2019) on videos with one, two, three, and four animals are 0.993,0.991,0.984, and 0.970, respectively. However, LMT only used one video to conduct MOTA calculations. Sample images and travel trajectories of mice can be observed in Fig 4C and in supplemental videos (4-7).

**Figure 4.**
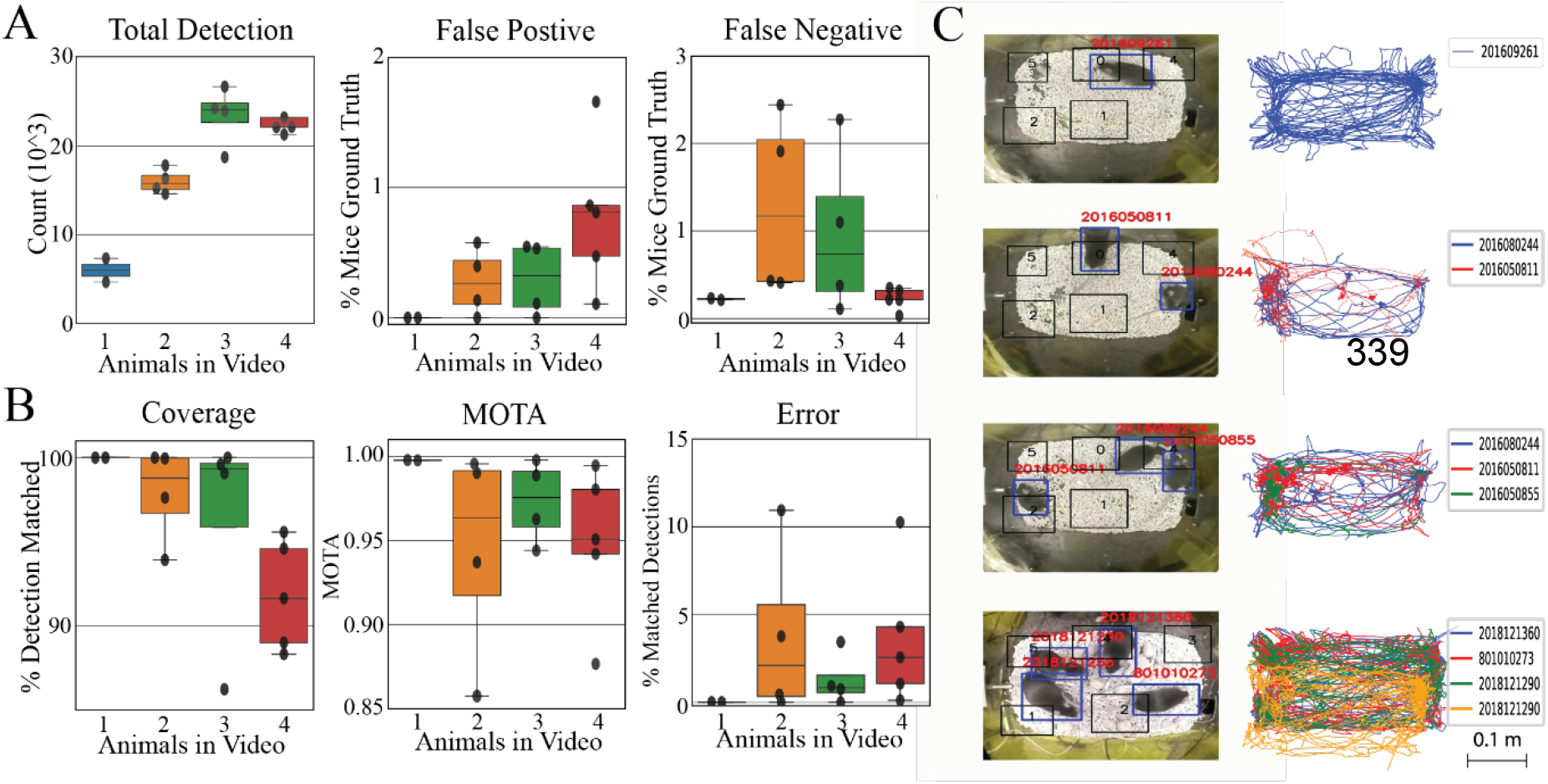
Evaluation of PRT performance in the modified home-cage. **A)** PRT detection Performance was measured by total detection, appearance of false negative detections, and false positive detections. False positive and false negative detections were expressed as a % of total ground truth mice in videos. Each individual data point represents a single 10-minute video **B)** Tracking performance of PRT was measured by coverage, MOTA, and % errors in detections with RFID matched. Coverage represents % of detections being matched with an RFID tag. As a reference for MOTA, LMT achieved values of 0.993,0.991,0.984,0.970 for one, two, three, four mice videos. Identity error is % of matched detection with an incorrect RFID tag. Each individual data point represents a single 10-minute video recorded in the home-cage. **C)** Sample image and travel trajectory of mice in the modified home-cage for cases with 1-4 mice.

### PRT Detection and Tracking Performance in Open-Field and Three Chamber Arena

To evaluate the scalability and customizability of the PRT tracker, additional behavior recordings were performed in an open field arena with dark bl6 black mice and in a three-chamber sociability arena with lighter coat color FVBN mice. In total, two videos were analyzed to evaluate the performance of PRT in an open field arena in a similar fashion illustrated in Fig 3B. In the two videos, FP and FN detections remain under 1 percent of total mice in ground truth (Supplemental Fig 2 and Supplemental Videos 8). Compared to PRT in the modified home-cage, the vast majority of detections can be matched with an RFID tag, with an average coverage of 99.99 percent between the two videos. Moreover, MOTA values were also very high in the two videos averaging at 0.97. Surprisingly, PRT achieved a very low identity error rate with that of both videos being less than 4%.

For the three-chamber arena, only one video was analyzed achieving similar performance in the open-field arena. After retraining Yolov4 on white mice detection, FN and FP were 0.03 and 0.14 percent respectively. MOTA index value was 0.965 and the identity error rate was 3.33 percent of all RFID matched detections. Sample illustrations of PRT tracking in a three-chamber arena can be observed in Fig 3C and in supplemental video 9.

### Stroke Induced Changes in Open-Field behavior

A total of 4 mice were recorded and used for analysis. Fig 5A illustrates a sample travel trajectory of a mouse pre-stroke, 1-day post stroke, and 7-day post stroke. Consistent with the literature, stroke acutely induces motor impairments in mice observable in the open field (Bains et al., 2018). A comparison of turn angle distributions of pre-stroke vs 1-day and pre-stroke vs 7-day post stroke shows a consistent pattern of decreased number of sharp angle turns and an increase of wide angle turns across the four animals tested as shown in Fig 5B indicating some recovery. A further analysis relieved statistical significant changes in distance traveled, mean speed, mean acceleration, number of sharp angle and wide angle turns not only between pre-stroke and 1-day post stroke (p<0.01), but also between 1-day post stroke and 7-day post stroke (p<0.05) in animals using a RM-ANOVA as shown in Fig 5C. Mean acceleration (p<0.05) and duration spent in center (p<0.01) were also different between pre-stroke and 1-Day post-stroke in animals.

**Figure 5.**
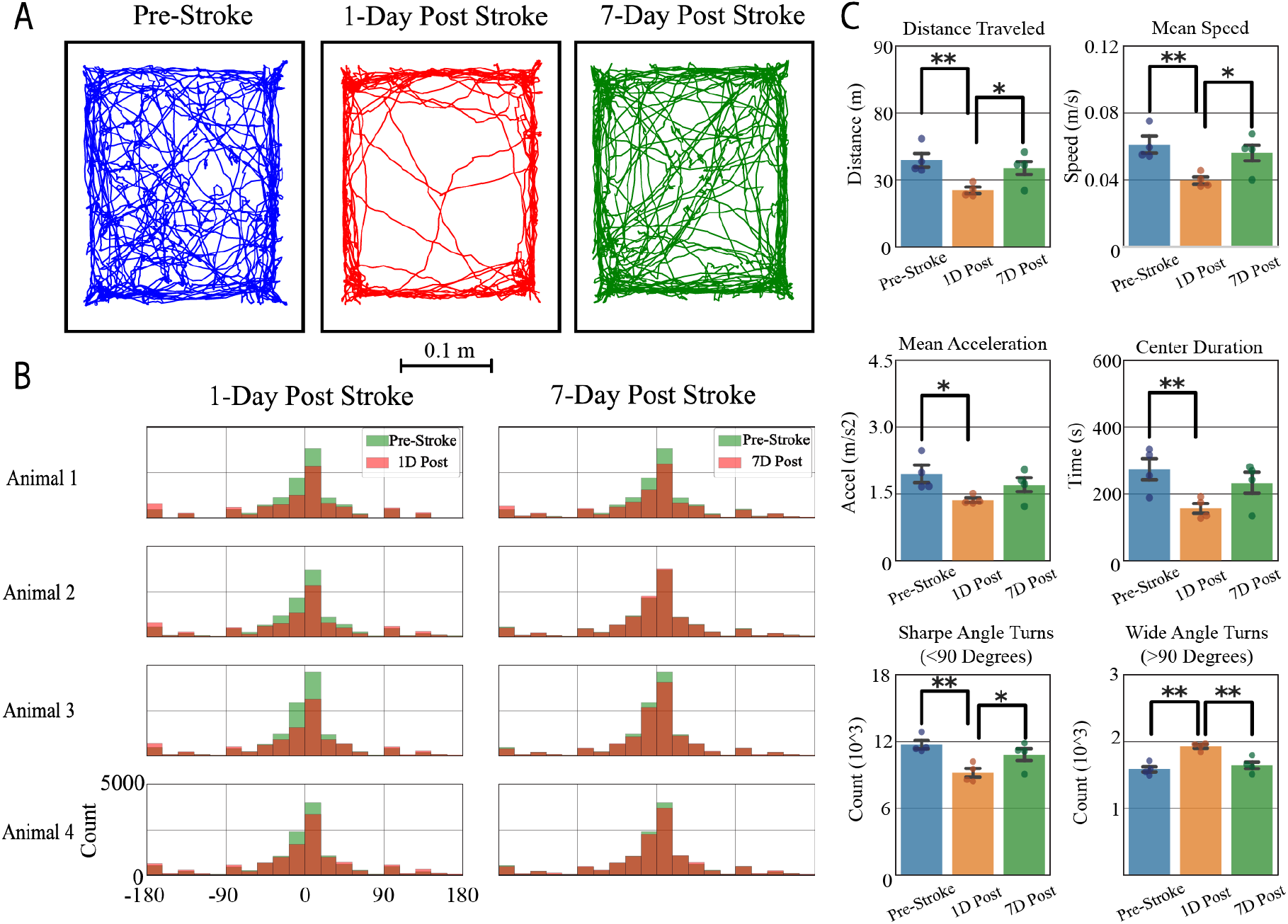
10-Minute Open-Field of Mice Pre-Stroke, 1 Day and 7 Days Post Stroke Induction. **A)** Sample travel trajectory of a mouse before, 1 day, and 7 days post stroke induction. **B)** Turn angle distribution of all stroke induced mice in open-field: pre-stroke vs 1 day post Stroke and pre-stroke vs 7 day post stroke. **C)** Significant changes in travel trajectory parameters detected by a one-way repeated ANOVA. * P<0.05 and ** P<0.01 (N=4 mice). Data are presented as the mean +/- standard error of the mean.

### Chronic Tracking in Custom Home-Cage

A total of 4 cages each containing 3 mice were recorded for three days using the modified home-cage illustrated in Fig 3A. The effects of day/night cycle (lights on and lights off) on dependent outcomes such as distance traveled, active duration, average speed, wide angle turn count, sharp angle turn count, and average acceleration along with anxiety measures such as duration spent in the center, interacting with one or two mice were determined using an MANOVA (Seibenhener and Wooten, 2015). As expected, mouse activity levels follow a reverse circadian rhythm (Ananthasubramaniam and Meijer, 2020).

Specifically, animals show higher levels of motor kinetics such as distance traveled (p<0.001; effect size=0.69), active duration (detected by the underlying motion detector; p<0.001, effect size = 0.54), average speed (p<0.01; effect size = 0.29), number of wide (>90 degrees; p<0.05; effect size = 0.2) and sharp (<90 degrees; p<0.001; effect size= 0.6) during periods of lights off as opposed to periods when the lights were on as shown in Fig 6A. However, no changes were observed in traditional anxiety related parameters such as duration spent in the center area, interacting with one or two mice as seen in Fig 6B.

**Figure 6.**
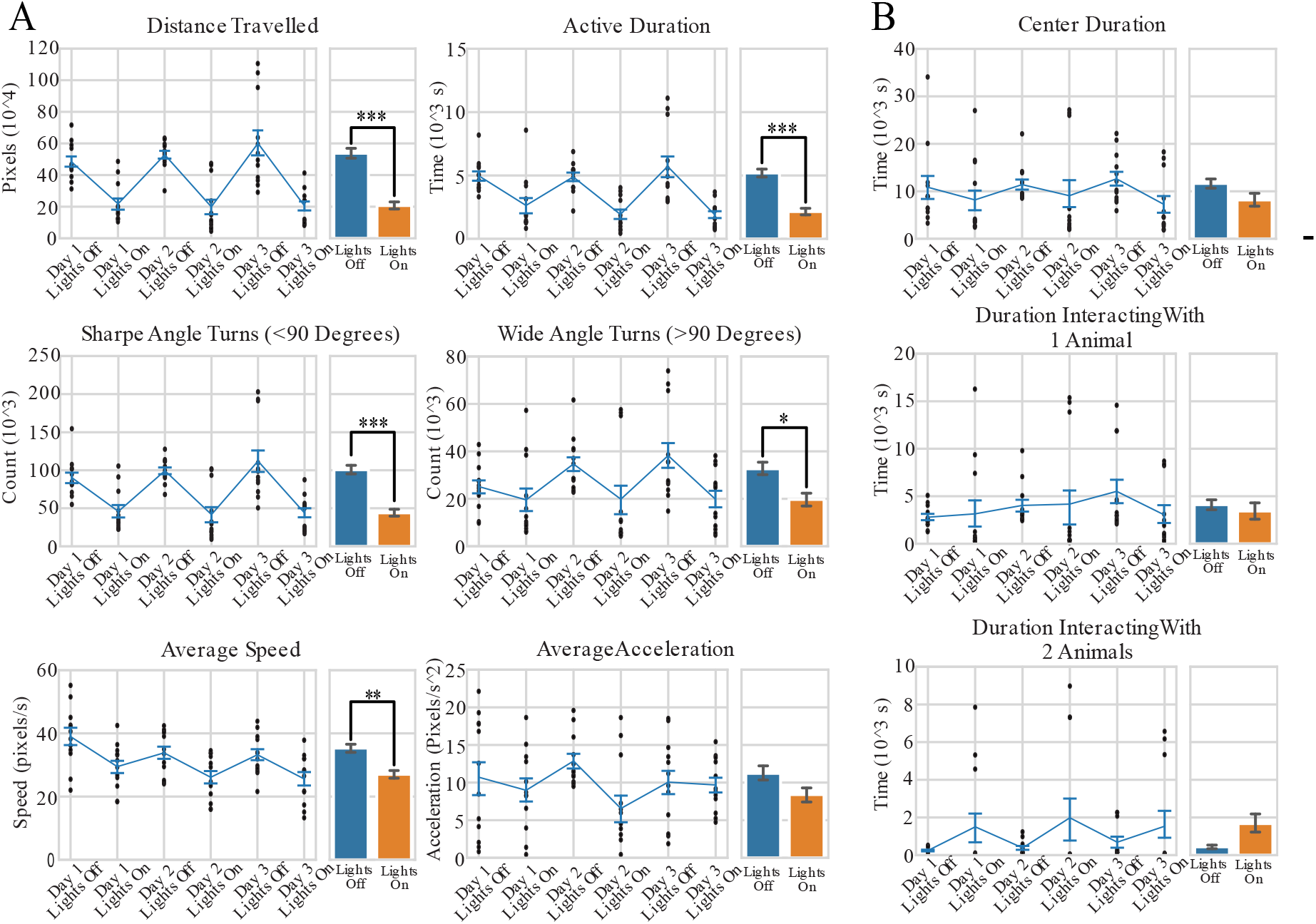
Analysis of 3-day behavior patterns in home-cage during day/night cycle (Lights on and lights off). **A)** Analysis of 3-day activity patterns. Total distance traveled by mice. p< 0.001; effect size = 0.69. Total duration of mice being active as detected by the motion detector. p<0.001; effect size =0.54. Number of sharp angle turns (<90 degrees) made by mice. p<0.001; effect size =0.60. Number of wide angle turns (>90 degrees) made by mice. p< 0.05; effect size = 0.20. Average speed of mice. p< 0.01; effect size= 0.29 **vi)** Average acceleration of mice detected. p>0.05. **B)** Analysis of 3-day activity patterns. Total duration spent in the center area of the cage, as defined by 0.5*width and 0.5*length around the center of the cage. No statistical significance was found in duration spent in the center, interacting with 1 mouse, nor interacting with 2 mice in daylight cycle. MANOVA was used with an ANOVA as post-hoc. Data= Mean +/- SEM (N=12). Each individual point represents a data point generated from a single mouse.

### Social Stimulus Test

Track pattern difference score, as its name implies, represents the similarity in pattern of the segmented pairs of travel trajectories regardless of length and spatial location. The larger the score would indicate a more dissimilarity in overall pattern between the trajectories. Therefore, comparison of identical trajectories would yield a value of 0. Spatial proximity is a measure of the distance/proximity between two trajectories, the lower the values, the closer the tracks are. Fig 7B and 7C illustrate sample trajectory comparisons of a single test mouse against a non-cagemate and a cagemate, respectively. During the 10 min duration of the recording, track pattern difference scores (p<0.05), total ITC duration (p<0.001), and number of ITC episodes (p<0.05) were found to be statistically significant whereas average duration per ITC was not (p>0.05) as seen in Fig 7C. For comparison to the more common 5 min of video recording used in the literature, the first 5 minutes was also separately analyzed as seen in Fig 7D which resulted in a similar pattern. Interestingly, the track pattern difference score is only statistically significant (p<0.05) when comparing proximal segment pairs (SP<300, proximal are potentially more similar) and not distal segment pairs (SP>300) as shown in Fig 7F shows all the segment pair comparison of an illustrative test animal to a cagemate and non-cagemate stimulus animal.

**Figure 7.**
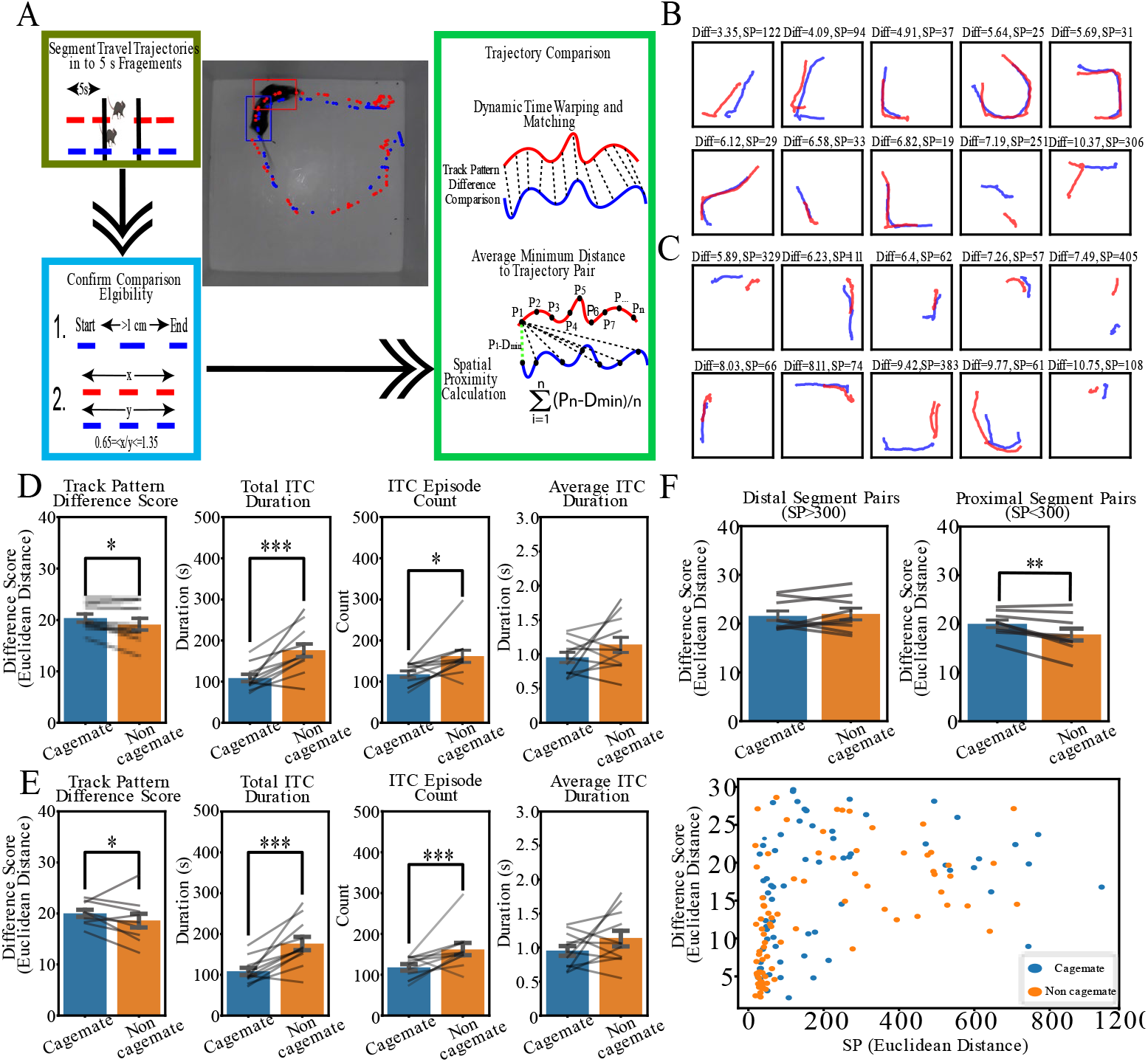
Travel trajectory analysis in an open-field social stimulus test. **A)** The overall analysis of travel trajectory comparison. Travel trajectories from two mice were segmented into 5 s fragments. Segment pairs generated need to both have start-end displacement of > 1 cm and to be similar in length in which one trajectory is not 35% longer or shorter than the other. If both criterias are met, the track pattern difference score and SP are generated. **B)** Sample segment pair (10 lowest track pattern difference scores) comparison from non-cagemates in a subject mouse**. C) S**ample segment pair (10 lowest track pattern difference scores) comparison from cagemates in a subject mouse. **D) S**ocial stimulus test result for 10 min. Average track pattern difference score, p<0.05. Total interaction duration, p<0.001. Number of social interaction episodes, p<0.05. Average duration of each social interaction episode, p>0.05. (N=11 mice) **E)** Social stimulus test result for the first 5 min of recording. Track pattern difference score, p<0.05. Total interaction duration, p<0.001. Number of social interaction episodes, p<0.001. Average duration of each social interaction episode, p>0.05. (N=11 mice) F) Track pattern difference score segregated by proximal (SP<300) and distal (SP>300) pairs. Track pattern difference score of distal segment pairs, p>0.05. Track pattern difference score of proximal segment pairs, p<0.01. (N=11 mice). Sample data in one test animal of all track pattern difference scores and SP values between segment pairs. Paired t test was Data= Mean +/- SEM . Each individual point represents a data point generated from a single subject mouse.

## 4. Discussion

In recent years, there have been a number of solutions to automatically capture the behavior and movements of specific rodents. These tools utilize technologies including combinations of computer vision, machine learning, neural networks, and physical tagging (Chaumont et al., 2019; Kiryk et al., 2020; Romero-Ferrero et al., 2019; Segalin et al., 2021; Singh et al., 2019; Walter and Couzin, 2021). In particular, video tracking coupled with radio frequency identification (RFID) has become a popular and reliable approach for identifying individual mice among groups without the use of visible markings (Bains et al., 2018; Chaumont et al., 2019; Peleh et al., 2019). A physical tagging such as RFID is necessary for complete automation of accurate tracking as each identity error in a crossing between mice can propagate throughout the rest of the video without a method of periodic re-verification (Branson, 2014). Although there are markeless trackers available for multi-animal tracking, these tools require manual validation/correction of animal identities (Lauer et al., 2021), animals to be of different coat colors (Segalin et al., 2021), or require videos to have uniform lighting and high animal contrast to background (Romero-Ferrero et al., 2019; Walter and Couzin, 2021). Hence, most current open-source automatic tracking methods are rather restrictive and in turn have only narrow application within particular arenas and experimental paradigms (Geuther et al., 2019).

Most automated tools, while desirable, remain difficult to access within the broader neuroscience community due to high barriers of costs, setup time, and ease of use. At the time of writing, there are three available systems that utilize video tracking coupled with RFID to capture mouse behavior in a home-cage like setting. Of these systems, two are commercially available: 1. RFID-Assisted SocialScan (Peleh et al., 2019) (CleverSys Inc., Reston, USA) and 2. Home-cage analyzer (Redfern et al., 2017). (Actual Analytics Ltd, Edinburgh, UK). Commercial tools are not only expensive to use at scale, but also cannot be modified to suit the investigator’s needs. Flexibility and scalability is especially important as investigators are employing more and more sophisticated approaches to tackle the increasingly complex questions being proposed. It is therefore of vital importance to be capable of tracking rodents of all coat colors and sizes in many different environments.

Additionally, software coupled with these commercial tools provides additional barriers for effective data and analysis method sharing.

As of writing, live mouse tracker (LMT) is currently the only open-source RFID and video tracking system available and is a milestone in multiple mice tracking (Chaumont et al., 2019). However, LMT has some limitations: 1) requires users to be experienced in configuring and setting up the corresponding hardware and software such as camera and RFID reader calibration (Shenk et al., 2020), 2) the setup is employed only in a defined large arena and does not seem adaptable/scalable to other recording environments such as a home-cage configurations, or the typically- sized mouse shoebox cages; and 3) installation of multiple systems will be more costly than our proposed system. LMT, while powerful, does require a desktop Windows computer which increases the potential footprint of the system and this could preclude its use at larger scales within an animal facility. According to the LMT build guide, the cost is approximately 3000 USD for one complete system. Although the associated costs of LMT are considerably lower than other commercially available solutions, it may also be a barrier to build multiple caged-based systems.

PRT offers multiple advantages over LMT: 1). affordability, 2) scalability, 3) ease of setup/use 4) customizability. In terms of cost, a PRT recording system with 6 RFID readers would cost approximately 520 USD, well below the 3000 USD of one complete LMT system. Affordability combined with PRT’s smaller footprint recording system would enable investigators to implement and use at larger scales, for example, monitoring racks of mice in an animal facility. In addition, the PRT recording system is very easy to set up and use. Aside from executing a setup script after connecting the RFID readers to RPi, no additional hardware calibration or installation is needed. Most importantly, the key distinction of PRT is the ability to be customized for tracking different types of rodents (varied coat color or species) in different experimental settings. As more and more in depth questions are being asked, the complexity of experimental designs have also increased (Basso et al., 2021; Slivkoff and Gallant, 2021). Investigators will need readily adaptable tools for their experimental design involving rodents of different size and coat colors in different environments. As shown in Fig 3(B-C), PRT can be adapted to record and track mice in a variety of environments, other than our home-cage configuration. In particular, PRT can be retrained to recognize mice of different colors even in cases where there is low mouse to environment contrast.

At the same time, PRT tracking performance of darker coat mice is slightly inferior to that of LMT (MOTA 0.97 for four mice) in exchange for the benefits mentioned above. Indeed, LMT can also further extrapolate postures and types of social interaction behaviors utilizing depth information by the use of depth sensing cameras. However, PRT also has the capability to incorporate output of posture estimation from multi-animal Deeplabcut which may enable similar features in the future (Supplemental Figure 3 and Video 10).

A substantial difference in PRT tracking performance was observed between our custom home-cage and more popular open arena settings. PRT performed higher in terms of detection and tracking in open arenas compared to our custom home-cage (19 x 29 x 12.7 cm). The performance differences observed were likely related to two factors: bedding and the existence of an entrance area. In the home-cage, bedding acted as a physical barrier increasing the distance between the RFID tag and RFID readers, in turn, decreasing the likelihood of an RFID tag being read when crossing a reader. From our group’s previous data, the read range of our current RFID reader using the current antenna (Sparkfun, ID- 20LA) was ∼30 mm (Bolaños et al., 2017). Given that bedding tends to clump (such as the common aspen wood shavings), this would further aggravate reader rangeerror from our experience. When using a home-cage setting similar to ours, we recommend a pellet type bedding which is less likely to clump and easier to shift around. In regards to the entrance area, it is a source of FN, FP, and identity error rate increases as it not only provides an area for mouse clumping, but also where mice can exit or enter the cage challenging detection with new IDs. Therefore, we recommend other investigators to take account these two factors when using PRT to design their own experimental paradigms.

While small homecages can present some challenges, accuracy is still high and PRT has not only demonstrated capabilities to chronically track multiple mice from minutes to days, but also detect underlying animal phenotypes (stroke). Consistent with other studies, chronic recordings in the modified home-cage revealed diurnal locomotor activity patterns in mice (Ananthasubramaniam and Meijer, 2020; Wang et al., 2019). Specifically, mice showed higher levels of distance traveled, turns made, travel speed, and duration being active at night compared to day cycles. In an open-field arena, mice showed locomotor deficits immediately after stroke induction similar to other studies (Shenk et al., 2020; Shvedova et al., 2021).

Stroke induced anxiety behavior (Larpthaveesarp et al., 2021; Vahid-Ansari et al., 2016) was also observed in mice as shown by less time spent in the center zone. Interestingly, it was observed that mice made a higher number of wide angle turns (>90 degrees) and lower number of sharp angle turns (<90 degrees) after stroke. A possible explanation for this is that high motor coordination between limbs is required to make specific turn angles. However, motor coordination is impared due to stroke (Pollak et al., 2012; Wang et al., 2018), which lead to unsuccessful sharp angle turn attempts resulting in wide angle turns being made instead. Further validation of turn angle parameters may yield an assessor of motor coordinations in openfield paradigms.

Deficits in social functions are key symptoms in many mental disorders including autism, depression, and schizophrenia (Benekareddy et al., 2018; Fujikawa et al., 2022). Currently, the most widely used pre-clinical test in rodents is the three chamber sociability and social preference for novelty assay for rodents (Golden et al., 2011; Kim et al., 2019). The main outcome of interest is the duration spent by the subject mouse in a chamber containing a stranger (non-cagemate) mouse compared to that in a chamber of a familiar (cagemate) mouse, both of which are confined by a wire cage. In general, a normal mouse would have a higher preference in investigating a stranger compared to a familiar mouse as most commonly measured by the duration spent together and number of approaches (Beery and Shambaugh, 2021; Golden et al., 2011; Kim et al., 2019). However, mouse social interactions are complex and affected by the states of both participating parties presenting the possibility that confinement of a mouse may risk reducing ecological validity. Here, PRT would allow the investigation of social preference in freely moving mice pairs. Indeed, mice still tend to have an increased duration of total interaction time and number of interactions with a stranger mouse compared to a familiar mouse.

Recent evidence has suggested that the travel pattern and actions of each individual animal may be influenced by the position and social status of another animal(Duvelle and Jeffery, 2018). Following this line of thought, we applied dynamic time warping, a method common in travel trajectory pattern comparisons (Brankovic et al., 2020), to compare similarities of trajectory patterns of a subject animal to that of a stranger or familiar mouse. Interestingly, it was found that travel trajectories of non-cagemate pairs tend to be similar when in spatial close proximity compared to that of cagemate pairs. Therefore, the comparison of travel trajectory patterns may offer a novel method in determining sociability of mice pairs that could provide an adjunct to the 3-chamber sociability test.

Overall, we have demonstrated the effectiveness of the PRT tracking system for tracking visually different rodents in a range of experimental settings and have shown the system’s ability to detect pathological changes in motor kinetics induced by stroke. We hope that this tool can enable the use of more complex experimental paradigms. In the future, we will also be uploading new detection weights of a wide variety of mice in different environments for users to directly use or as bases for transfer learning to train their own weights.

---

**Supplementary Figure 1.**
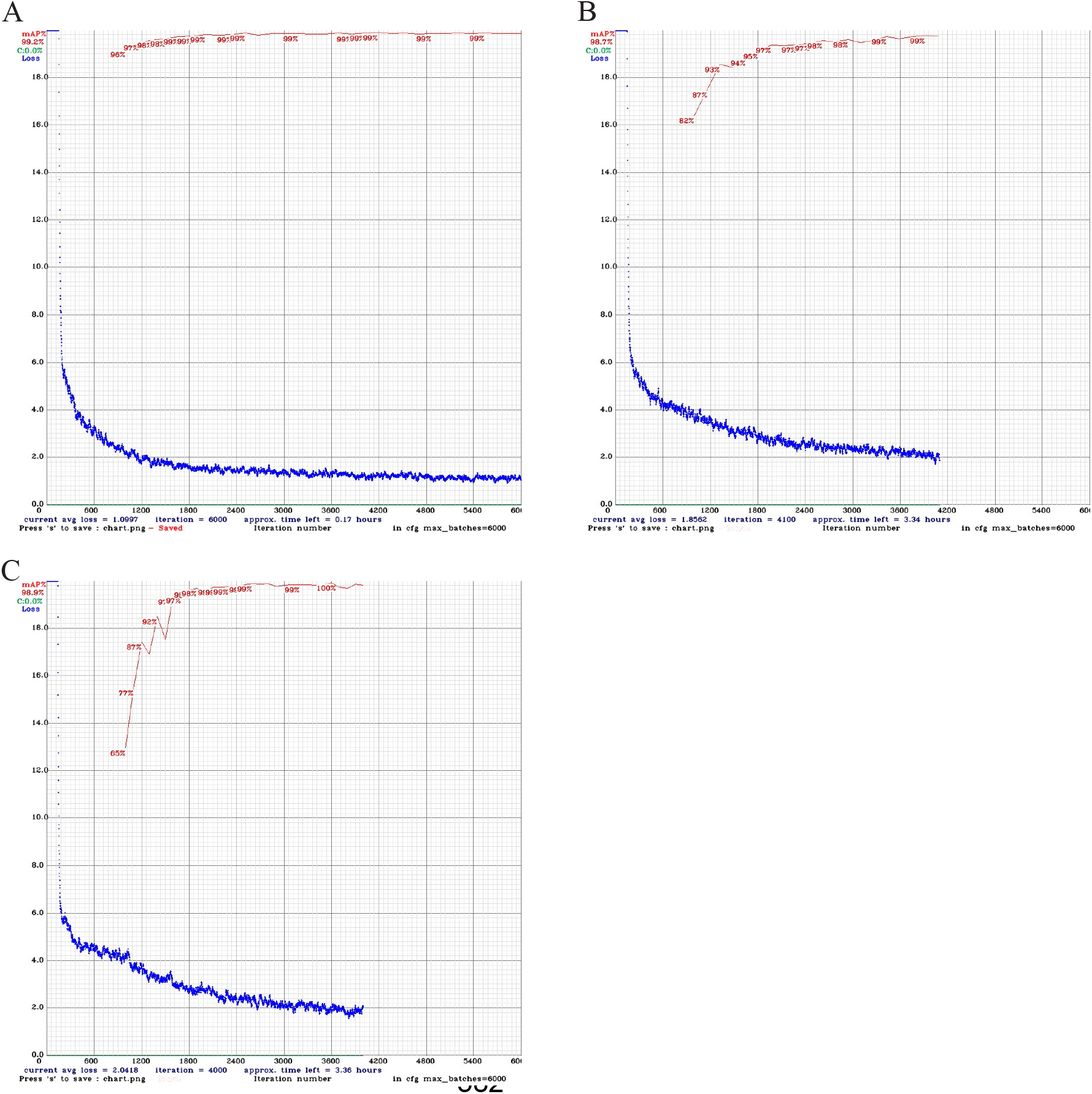
Yolov4 Training Loss and mAP. A) Training loss and mAP on home-cage weights using 2000 images with 6000 iterations. B) Training loss and mAP on the open-field arena using 300 images with 6000 iterations. Training was manually stopped when mAP > 99.5 %. C) Training loss and mAP on the sociability chamber arena using 300 images with 6000 iterations. Training was manually stopped when mAP > 99.5 %

**Supplementary Figure 2.**
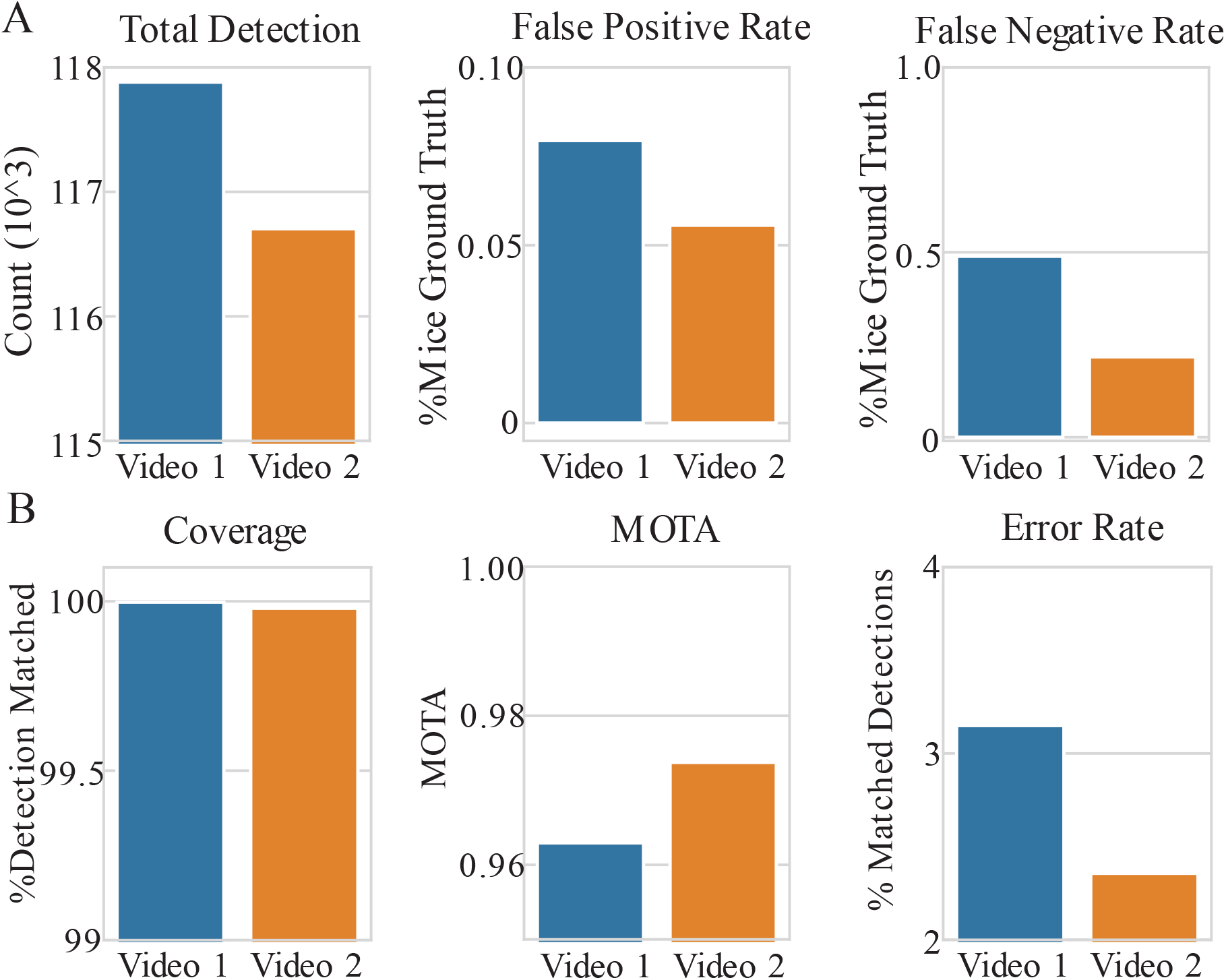
Evaluation of PRT performance on the openfield arena with 6 mice. **A)** PRT detection Performance was measured by total detection, appearance of false negative detections, and false positive detections. False positive and false negative detections were expressed as a % of total ground truth mice in videos. **B)** Tracking performance of PRT was measured by coverage, MOTA, and % errors in detections with RFID matched. Coverage represents % of detections being matched with an RFID tag. A total of two videos were evaluated.

**Supplementary Figure 3.**
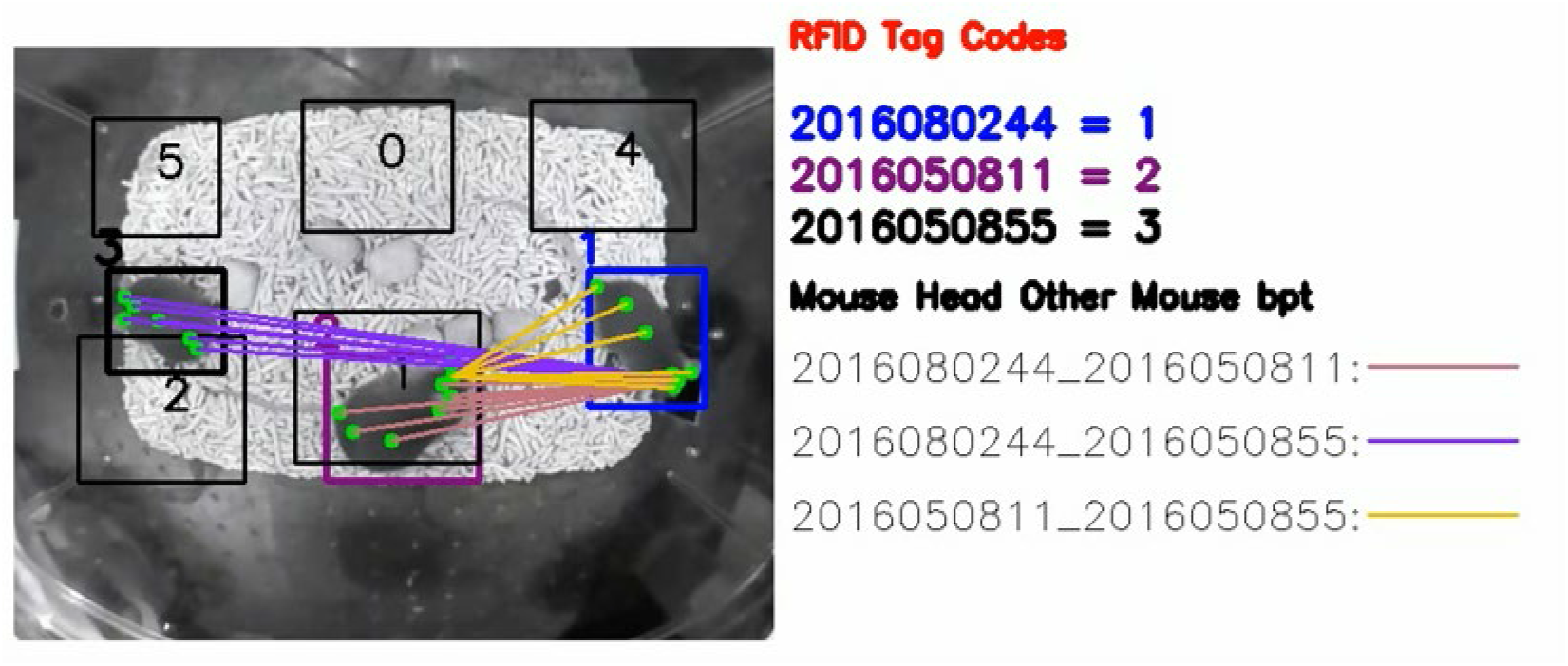
PRT with Deeplabcut output. Incorporation of Deeplabcut posture estimation with PRT identity tracking. Body parts for each animal tracked include snout, head center, left ear, right ear, neck, mid body, lower mid body and tail base. Distance of each mouse’s mouse head to other mice’s body parts are also shown.

**<u>Supplemental Video 1</u>. PRT software setup on a RPi.** A detailed walkthrough of setting up PRT online recording system on a RPi. This tutorial is also intended for users with a more limited background in coding or use of Linux-based systems.

**<u>Supplemental Video 2</u>. PRT demo hardware setup.** A demonstration of setting up the PRT in an openfield arena with 9 RFID readers. The total duration of the setup is less than 30 minutes. Extra tools include M1 screws and a glue gun.

**<u>Supplemental Video 3</u>. Modified SORT algorithm tracking during high occlusion and clustering situations.** A demonstration of the modified SORT algorithm tracking mice during scenarios of occlusions and mice clumping/clustering together. With Yolov4 and the original SORT algorithm, many detections are lost.

**<u>Supplemental Video 4</u>. PRT Tracking of one mouse in custom-built home-cage.** Each video frame is recorded at resolution of 512 x 400 at 15 frames per second.

**<u>Supplemental Video 5</u>. PRT Tracking of two mice in custom-built home-cage.** Each video frame is recorded at resolution of 512 x 400 at 15 frames per second.

**<u>Supplemental Video 6</u>. PRT Tracking of three mice in custom-built home-cage.** Each video frame is recorded at resolution of 512 x 400 at 15 frames per second.

**<u>Supplemental Video 7</u>. PRT Tracking of four mice in custom-built home-cage.** Each video frame is recorded at resolution of 512 x 400 at 15 frames per second.

**<u>Supplemental Video 8</u>. PRT Tracking of six mice in an openfield with 9 RFID readers.** Each video frame is recorded at a resolution of 960 x 960 at 40 frames per second.

**<u>Supplemental Video 9</u>. PRT Tracking of three white color coated mice in a three chamber arena.** Each video frame is recorded at a resolution of 640 x 480 at 40 frames per second.

**<u>Supplemental Video 10</u>. PRT Tracking with Deeplabcut posture estimation.** Body parts for each animal tracked include snout, head center, left ear, right ear, neck, mid body, lower mid body and tail base.

## Custom Home-Cage Parts

All related CAD files can be found at our github: https://github.com/tf4ong/tracker_rpi.

## 3D Printed Parts (Black PLA)

RFID_reader_base.stl

Nest_tunnel.stl

Nest_body.stl Nest_lid.stl

Camera_LED_mount.stl

Cage_Lid.stl

## Data availability

All data will be uploaded to the OSF data repository: https://osf.io/78akz/?view_only=c5d81b6162564249b8ce128d4d34abe1

## Acknowledgments

This work was supported by Canadian Institutes of Health Research (CIHR) Foundation Grant FDN-143209 to T.H.M. and the UBC Institute of Mental Health Marshall Scholars and Fellowship program. THM is supported by the Brain Canada Neurophotonics Platform and the Canadian Partnership for Stroke Recovery. We thank Pumin Wang and Cindy Jiang for surgical assistance and Jeffrey M LeDue for technical assistance. We thank Pankaj Gupta for the use of stroke and control group mice. This work was supported by resources made available through the Dynamic Brain Circuits cluster and the NeuroImaging and NeuroComputation Centre at the UBC Djavad Mowafaghian Centre for Brain Health (RRID SCR_019086) and made use of the DataBinge forum.

